# Accurately Programming Complex Light Regimes with Multi-channel LEDs

**DOI:** 10.1101/2025.06.06.658293

**Authors:** Gina Y.W. Vong, Paul Scott, Will Claydon, Jason Daff, Katherine Denby, Daphne Ezer

## Abstract

**Background:** Advances in LED lighting technologies have allowed researchers to explore increasingly complex light regimes. This has given us greater insight into plants’ responses to dynamic light, including seasonality and fluctuating conditions, rather than the discrete (i.e. on / off) lighting previously explored. However, there is a need for methods to accurately program multi-channel / waveband LED lighting systems.

**Results:** We present a multi-step, multidimensional algorithm to accurately program LED lights. This algorithm accounts for non-linearity between intensity settings and irradiance output, as well as bleedthrough between channels of different wavebands. Our algorithm out-performs other methods which treat waveband channels as independent variables, more accurately predicting intensity settings to achieve a desired irradiance when using multiple LED channels.

**Conclusions:** This algorithm allows the community to accurately program complex light regimes to probe plant responses to dynamically changing light spectra. We have made this algorithm available to the plant science community as an R package, LightFitR (available on GitHub at: https://github.com/ginavong/LightFitR).

## Introduction

Plants rely on light to fix carbon and grow. As such, light is an essential source of information for plants [1] to optimise their photosynthesis. Red / far-red ratios drive shade-avoidance [2–4], while blue light is involved in photomorphogenesis [5–8]. Plants have thus evolved elaborate mechanisms for gathering this information, integrating it with other environmental inputs and translating it into a physiological response [9–12]. Indeed, much work has been done to study the mechanisms of how plants perceive and respond to light information. These insights allow us to understand, with increasing resolution, how a plant behaves, with the potential to inform strategies to engineer climate resilient crops and novel agricultural technologies [13,14].

Light information can be subdivided into 3 components: i) irradiance, how much light the plant is receiving [15]; ii) photoperiod [16], the duration for which the plant is receiving light; and iii) light quality, corresponding to the spectrum and wavelengths / colours of light [17]. Together, these 3 components allow the plant to infer information about the time of day [18], seasons [19–21] and shade avoidance [2].

However, much of the work on plant light response to date uses discrete lighting changes (e.g. sudden on/off) or constant light/darkness [9,10], rather than the dynamic changes seen in nature where irradiance, photoperiod and spectrum often gradually change over the day and across seasons [22]. Increasingly, research has focussed on plant responses to fluctuating light conditions throughout the day [11,12,23–27], giving us a deeper understanding of plants’ perception and response to light signals, as well as implications of variable light on plant productivity. These studies take advantage of advances in LED technologies [28,29], which allow researchers to modulate the irradiance across multiple waveband channels in pre-defined schedules to produce dynamic light regimes. However, there is currently inconsistent reporting of lighting regimes used in these experiments [30], and a lack of methods for accurately programming complex light regimes involving multiple channels at different irradiances.

Existing methods for calibrating and programming LED lights vary. Some use regression methods [31] while others seem to use a laborious ‘measure and adjust’ approach [32]. Regression-based methods assume a linear relationship between intensity (settings within the LED, typically a %) and irradiance (measured light output from the LED). Further, by treating each LED channel individually (i.e. assuming independence between the channels), neither approach accounts for the complex interaction of the spectral irradiances produced by multiple overlapping channels. A spectrometer (rather than a light meter) allows the assessment of the wavelength range of each LED channel, the extent of overlap between channels, and whether a channel has a secondary peak [30,33]. All these factors can influence the output of the LED fixtures, as well as plant responses in experiments [30].

Moreover, the output of the lights is highly dependent on their installation, as different materials on growth cabinet walls have differing reflectance and absorbance. This affects how much, and which wavelengths of, light actually reach the plant, not to mention the potential variation between LED fixtures even of the same model. Therefore, each LED lighting setup requires a bespoke calibration in order to accurately program a light regime for experiments.

Here, we present an experimental protocol coupled with an algorithm for effectively calibrating and programming complex light regimes involving multiple LED waveband channels across a range of irradiances. We also present a further refinement step for those requiring additional accuracy. The algorithm is available as an R package, LightFitR, facilitating the employment of more complex light regimes for plant science experiments.

## Methods

All data processing, analysis and figure creation was carried out in R (version 4.4.1). The code is available on GitHub at: https://github.com/ginavong/2024_LightFitR_MethodsPaper and the raw data is available at: https://doi.org/10.5281/zenodo.15584172

### Defining key terms

In the context of this manuscript, we use the term *LED channel* to denote the waveband produced by a given type of LED. While *LED fixture(s)* refers to the lighting fixture as a whole, including all the LED channels housed within.

We use the term *intensity* to refer to the unitless setting we provide to each LED channel. In contrast, we use the term *irradiance* to refer to the instantaneous measurement (area x time) of light output that is present at specific designated wavelengths (in nm) at the plant level in the growth chamber. The aim of our protocol is to enable a researcher to predict what intensity they should programme for each LED channel to produce a *target irradiance* at a target waveband required for a specific experiment.

*Peak wavelength* is the precise wavelength where the maximum irradiance is observed for that channel. We define *bleedthrough* as the irradiance of light from one LED channel, which is encroaching on the peak wavelengths of other channels.

Further, we use *event* to refer to the time period when a given set of LED channels is on at a given set of intensities. A series of events is referred to as a *regime* or *schedule*.

### Cabinet setup

The experiments were conducted in a Snijders (Tilburg, Netherlands) MC1750 growth chamber with Internal dimensions (w x d x h) 1830mm x 780mm x 1230mm. The chamber was modified to provide two equal growing compartments with separate lighting regimes, by installing an internal partition made from white Palfoam (Palram, Hull, UK) aerated PVC foam sheet. The partition was sealed with RS PRO (Corby, UK) 5mm Black Foam Tape, to prevent light spillage between the compartments and did not impair the upward airflow distribution system (Fig.1A). Sufficient outdoor make-up air provides ambient CO2 conditions inside the chamber. The chamber air temperature was maintained at 22/22°C (s.d. ±1/1°C) during the light/dark period. The relative humidity in the chamber was maintained at 55% (s.d. ±10%).

**Figure 1.**
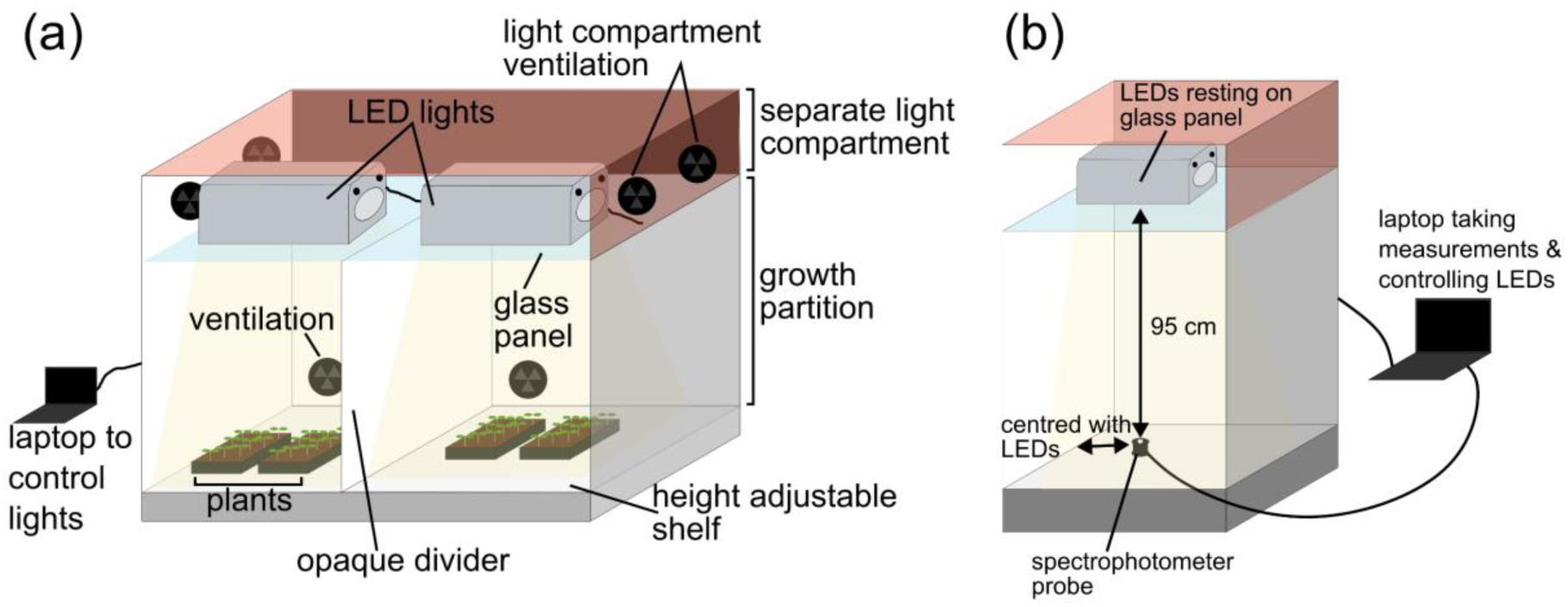
Cabinet setup with Heliospectra DYNA lights. **(a)** Overall setup of the growth cabinet with 2 Heliospectra DYNA lights resting on a glass shelf in a separate lighting compartment with its own ventilation. These lights are connected to a laptop which controls them. The growth compartment is separately ventilated, with a height adjustable shelf for plants. An opaque divider was installed to separate the light coming from the 2 sets of LEDs above. **(b)** Setup for taking measurements. Only half of the cabinet is shown. The spectrometer rests on the shelf, 95cm away from the light source. The spectrometer is connected to a laptop which records the measurements while also controlling the LED lights.

Heliospectra DYNA (Gothenburg, Sweden) LED fixtures were mounted in the existing passively ventilated lamp loft, partitioned by a clear glass barrier above the growing compartments below, the LED fixtures were cooled by internal variable speed fans. The LED channels were controlled by helioCORE™ (R3.2.2-Release) a web-based lighting control system, independent of the growth chamber’s local control system.

### Light Measurements

For automated light measurements, schedules were uploaded to the LED fixtures via helioCORE. Each event in the schedule lasted 5 minutes to allow time for multiple readings to be taken.

Light readings were taken at the plant growth level, 95 cm below the centre of the LED fixture (Fig. 1B). For spectrum readings, an OceanInsight FLAME spectrometer (Ocean Optics, Duiven, Netherlands) was used in conjunction with OceanView (version 2.0.14) software recording every 1 minute with a 500 ms integration time and averaging over 5 scans.

### Spectrometry data processing

The spectrum measurements are processed to trim the wavelengths from 300-800 nm. Irradiance unit conversions (from μW cm^-2^ nm^-1^ s^-1^ to μmol m^-2^ nm^-1^ s^-1^) were calculated using the photobiology package (version 0.11.3) in R. As there were multiple readings for each event, the reading at the middle time point was selected for subsequent analysis. Further annotation of the processed dataset was then carried out depending on the analysis required (see specifics below).

### Generating and analysing calibration data

Calibration curves were generated separately for each LED channel (380 nm, 400 nm, 420 nm, 450 nm, 530 nm, 620 nm, 660 nm, 735 nm and 5700k white), by increasing the intensity every 5 minutes (0, 1, 5, 10, 20, 50, 100, 200, 300, 400, 500, 600, 700, 800, 900, 1000) and taking spectrometer readings every 1 minute (see ***Measurements of the Lights***).

After the raw measurements were processed (see **Spectrometry data processing**), the following values were calculated for each reading (Fig. 2A): (i) the total irradiance (i.e. sum of irradiances between 300-800 nm); (ii) the peak wavelength (i.e. wavelength at which the maximum irradiance occurred in the spectrum), with the median peak wavelength for the LED channel used for future analysis); and (iii) the irradiance at the peak wavelength (i.e. peak irradiance). Further, we calculated *bleedthrough* for each channel at 1000 intensity by extracting the irradiances at the peak wavelengths of the channels which were not on.

**Figure 2.**
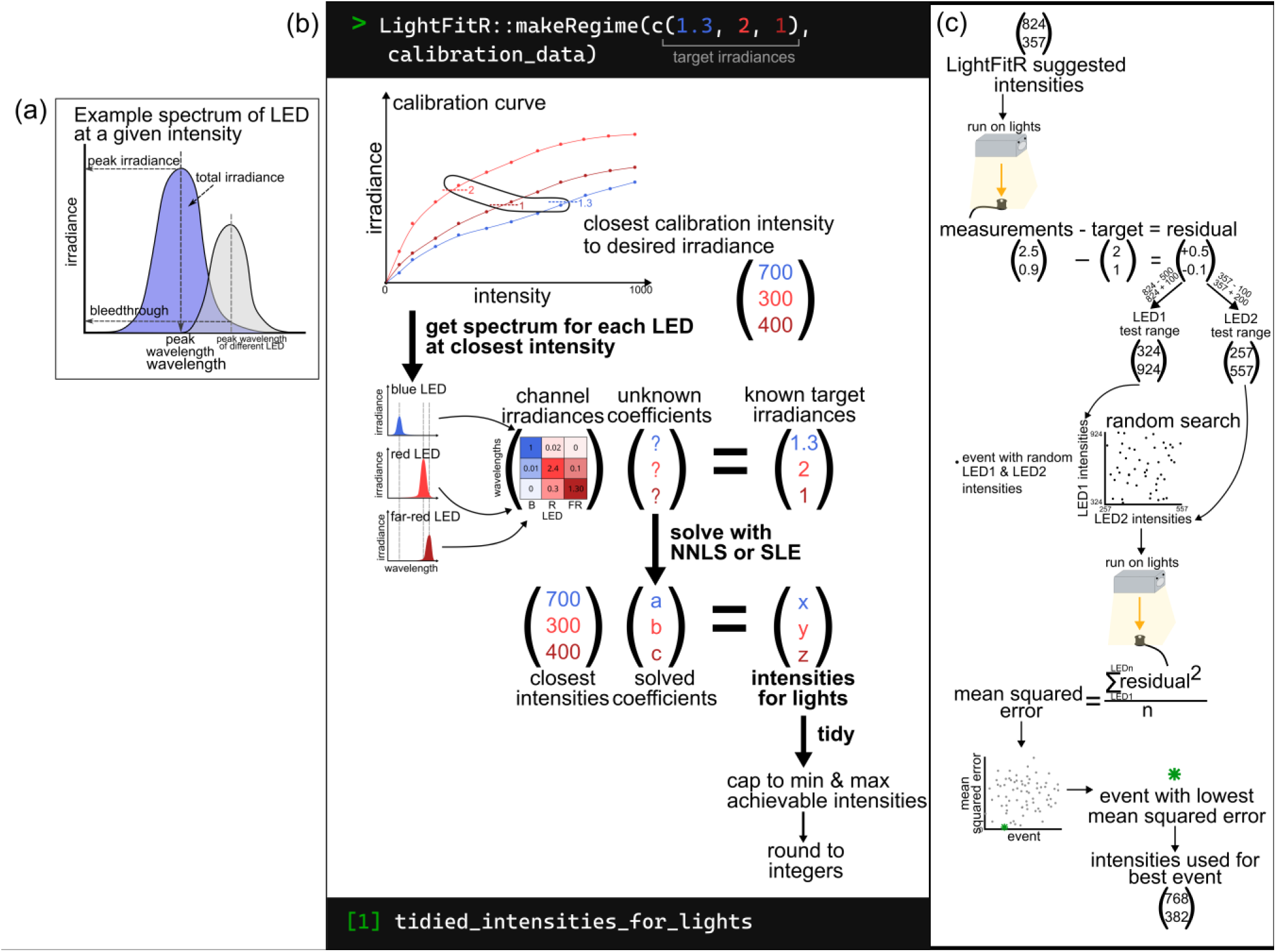
Analysis and algorithm diagrams. **(a)** Example spectrum from an LED at a given intensity, breaking down which features were used for analysis: peak wavelength, peak irradiance, total irradiance, bleedthrough. **(b)** Graphical explanation of our algorithm for accurately programming LED lights. It takes user provided target irradiances and calibration data and uses it to predict which intensities to set the lights, in order to achieve the target irradiances. This is achieved through multiple steps: obtaining the closest calibration intensity to the target irradiance, getting the calibration spectrum for each LED at that closest intensity, compiling a matrix of these channel irradiances, solving a system of linear equations with this matrix and the target irradiances, multiplying the solved coefficients with the closest intensities to predict the optimum intensities for the lights, and tidying the predicted intensities so that they are accepted by the lighting software. **(c)** The process of carrying out a random search. Take the intensities that the LightFitR algorithm predicts and run it on the lights, measuring their output. Calculate residuals by comparing the measurements with the original target irradiances. Test ranges for each LED are calculated proportional to the scale and direction (positive or negative) of the residual, using the LightFitR intensities. Random combinations of intensities (within these ranges) are selected and then run on the lights with each random combination considered an event. Measurements are again taken and mean squared error for each event is calculated using the residuals. The event with the lowest mean squared error is the ‘best’ event and the intensities used for that best event should be used in experiments.

### Intensity Prediction Algorithm

Our algorithm (Fig. 2B) takes user-provided calibration data and user-defined target irradiances for each peak wavelength in order to predict intensities for each channel. These intensities can be programmed into the fixtures to achieve the target regime. It consists of 3 steps: (i) produce a rough estimate of intensities by directly using the calibration data for each LED channel, (ii) adjust the estimated intensities to account for bleedthrough between LED channels, and (iii) modify estimated intensities to fit hardware requirements (Fig. 2B).

First, when given an event of target irradiances at the peak wavelengths, it searches the calibration data for intensities which produce the closest irradiances to the target irradiance for each channel (i.e. intensity with the smallest absolute difference in irradiance, called the *closest intensities*). We define *m* to be the vector of closest intensities for each LED channel.

Next, we refine these intensities to account for bleedthrough between channels (see ***Generating and processing calibration dat*a**). We combine bleedthrough at the closest intensities identified in the first step into a matrix, Mm.

This matrix is then used to solve a multidimensional system of linear equations, either using R’s ‘solve’ function (we simply call this approach ‘system of linear equations (SLE)’) or with non-negative least squares (NNLS, [34]), as in Equation 1. To calculate the predicted intensities, we use Equation (2).

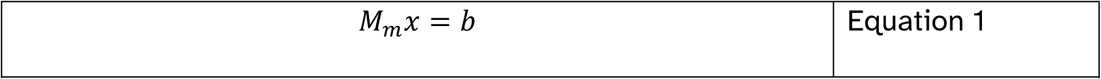

Where M is the bleedthrough matrix at LED intensities m, x is the vector of coefficients we are trying to predict and b is the vector of target irradiances.

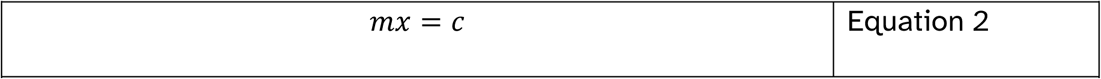

Where c is the predicted intensities that we expect will produce irradiances close to the target irradiance b.

Finally, we must adapt these estimates to meet the hardware requirements of the LEDs. For example, these predicted intensities may be below 0 (in the case of SLE) or above the maximum achievable intensity by the LEDs (1000 for Heliospectra DYNA). Furthermore, the LED fixtures can only take integer (whole numbers) intensity values while the predicted intensities are often not integers. Therefore, we created a final ‘tidying’ step which (i) rounds to the nearest integer; (ii) sets negative predicted intensities to 0; and (iii) caps the maximum intensity to the maximum achievable by the light model.

### Algorithm comparison

We compared our multidimensional algorithm (with SLE and NNLS) with other methods: individual closest, individual linear model / regression, individual NNLS.

We first designed a test regime where random LED channels were turned on in combination, at random intensities. Our testing regime increased in complexity, starting with just 2 random channels active / on at a given time and increasing until all 8 LED channels were on simultaneously, with n=20 events for each level of complexity. We ran this regime and measured the spectra for each event (see <u>light measurements</u> and <u>spectr</u>ometry data processing).

Using the measured irradiances, we asked each algorithm to predict the intensities which were originally used. This allowed us to assess the accuracy of the algorithm predictions for each LED channel at each event (i.e. the residuals: predicted - true). To produce an overall score for how accurate each algorithm was for a given event, the mean squared error (MSE) across LED channels was calculated for each event. We compared the algorithms at each complexity level using a Kruskal-Wallace test, with Dunn post-hoc test and Holm’s p-value adjustment. To assess whether an algorithm was consistently over- or under-shooting, we created visualisations of the raw differences/residuals.

### Refinement with random search

Additionally, we demonstrate experiments for further refining the intensities to produce irradiances that are even closer to the target irradiance, by using a random search approach (Fig. 2C). The premise is that we select random combinations of intensities near our algorithm’s predicted optimal intensities, which can be experimentally tested on the LEDs to enable further refinements. The range of the intensities is weighted by how ‘wrong’ the algorithm predicted intensities are.

More specifically, we programme the algorithm’s predicted intensities in our LEDs (i.e. the starting intensity), measure the irradiance at each waveband and calculate the difference between the actual irradiance and the target irradiance (i.e. the residual). We defined test ranges for each LED channel, which were proportional to the direction and size of the residual for the algorithm predicted intensity (see Equations 3 & 4). n=50 events then were selected for each treatment, each with a random combination of intensities (within the LED channels’ test ranges). These events were tested on the LEDs and the spectra collected. The optimal intensities are then defined as the combination of intensities which have the lowest MSE.

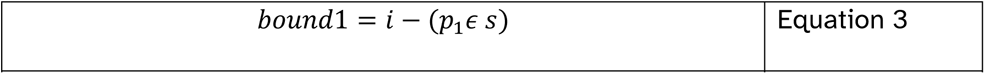

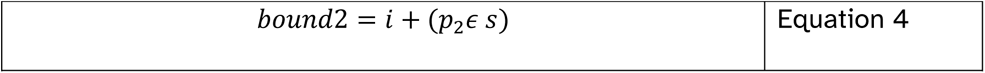

Where *bound1* defines an intensity limit in the opposite direction of the residual (*ϵ*) and *bound2* defines an intensity limit in the same direction as the residual. *i* is the algorithm predicted intensity, and *s* is a scaling factor turn the small residuals (in µmol s^-1^ m^-2^) into numbers >1 (we used s=100 but other setups may differ). *p* is a proportionality factor that indicates how far away from the algorithm predicted intensity should be searched where *p1 > p2* to ensure that the range of *bound1* is greater than *bound2* (we used *p_1_*=10^1.4^ and *p_2_*=10).

### Testing the efficacy of random search

We tested the random search approach for three different sets of target irradiances to use as ‘treatments’. This allowed us to assess the marginal benefit of using random search for different starting (i.e. algorithm predicted) MSEs. To determine the treatments, we used target irradiances from the ***Algorithm comparison*** section at a complexity of 4 LED channels, selecting the events with the lowest (minimum), median and highest (maximum) predicted MSE.

After processing the raw spectrometer data, the dataset was filtered to retain only the irradiances at the median peak wavelengths for further analysis. These were compared with the target irradiances to calculate residuals and then MSE per event. Within each treatment, the event with the lowest MSE was labelled the ‘best’. The Euclidean distance between each event and the corresponding ‘best’ event was then calculated, and Spearman rank used to assess the correlation between MSE and Euclidean distance.

## Results

### Properties of the lighting system

‘Intensity’ is an arbitrary unit within the light system settings, associated with the amount of electrical power that a particular channel receives. Thus we investigated how intensity impacts the amount of light (irradiance, μmol m^-2^ nm^-1^ s^-1^) being output. We noted that irradiance does not scale linearly with intensity (Fig. 3A & Fig. S1A). Each LED channel has a different maximum irradiance, with the 380 nm channel emitting particularly low irradiances of light. Moreover, the maximum irradiance for the 620 nm channel was around 600 intensity rather than 1000.

**Figure 3.**
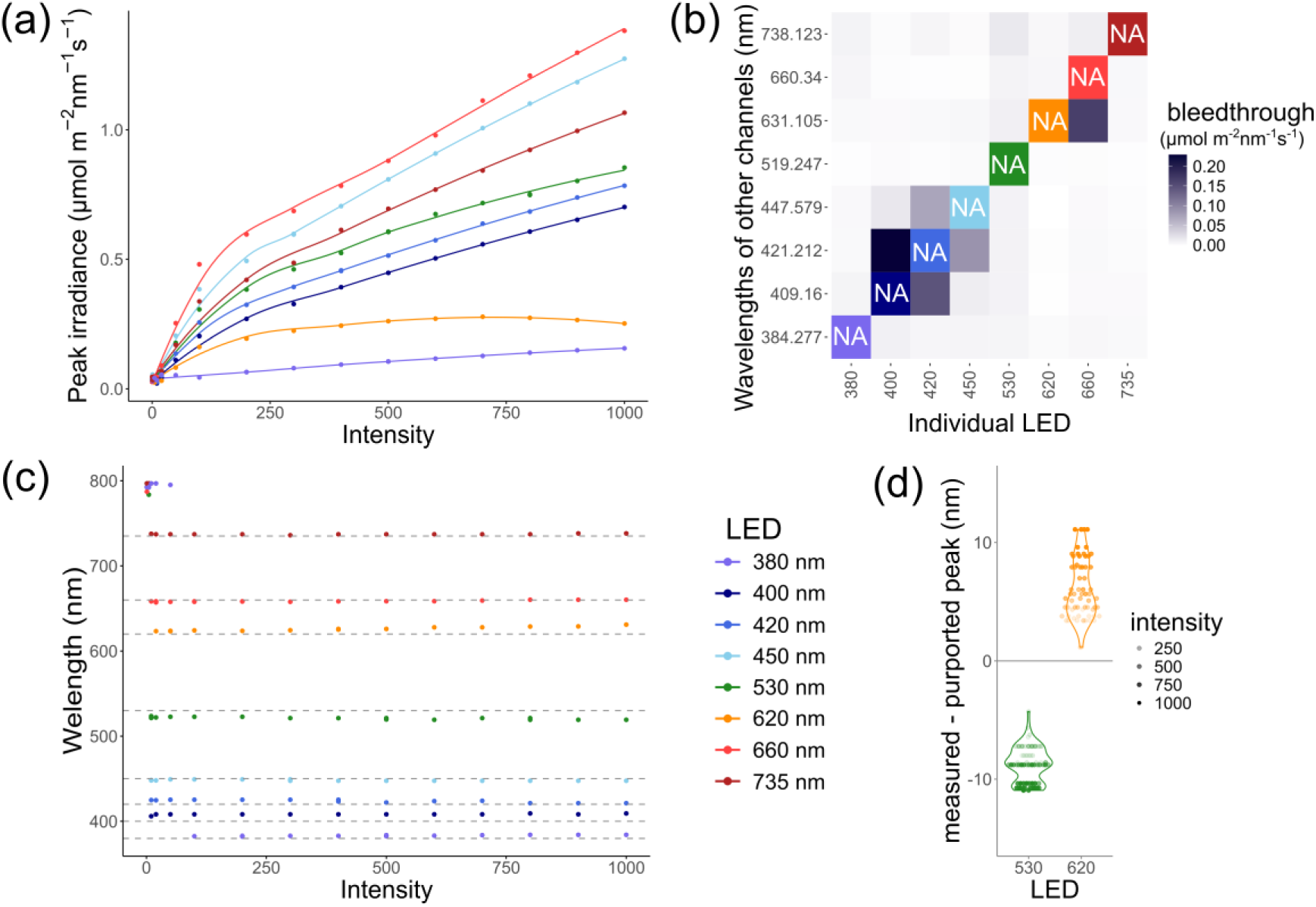
Challenges of the lighting system, revealed during calibration. The spectrum was measured independently for each LED channel at increasing intensities (0, 1, 5, 10, 20, 50, 100, 200, 300, 400, 500, 600, 700, 800, 900, 1000) with a spectrometer. **(a)** The irradiances at the peak wavelengths of the spectrum, with a smoothing function applied to highlight the non-linearity. **(b)** The bleedthrough from 1 LED into the wavelengths of the other channels, at maximum (1000) intensity. The diagonal is NA as that represents the channel which is on. **(c)** The wavelength of the spectrum peak for each LED channel, with dashed lines representing the purported peak wavelengths. **(d)** The difference between the peak we measured and the purported peak, and how it relates to intensity. The 530 nm and 620 nm channels are highlighted as they show the most extreme differences.

We also noticed that the spectra of some LED channels overlapped with each other (Fig.3B and S1B), resulting in bleedthrough. This bleedthrough is particularly severe among the blue (400 nm, 420 nm, 450 nm) channels, with the green (530 nm) channel bleeding through into all other channels and bleedthrough also evident between the red (620 nm, 660 nm, 730 nm) channels. We note also, a small amount of bleedthrough between the blue and red channels, and vice versa. The algorithm we developed accounts for both the non-linear associations between intensity and irradiance and the bleedthrough between LED channels.

Finally, we wished to confirm whether the peak wavelength for each channel matched the named wavelength of the channel (Fig. 3C). We note that none of the channel peaks exactly matched the purported wavelengths, with some channels deviating more than others. At low intensities, the noise from the spectrometer was greater than the irradiance at the peak wavelength, and thus the ‘peaks’ around 800 nm at those low intensities do not represent true output from the light. However, we noted that, at intensities over 100, the peak wavelengths were changing and diverging from the purported wavelengths as the intensity increased (Fig. 3D). Going forward, we define the peak wavelength for a channel as the median of that channel’s peak wavelengths across all intensities.

### Performance of the prediction algorithm

Our algorithm aims to accurately predict intensities for LED lights to achieve user- defined target irradiances and wavelength profiles. To test the performance of our algorithm, we took detailed measurements from a complex light regime which had random combinations of LED channels using the light set up described above. Using these measurements (irradiance and wavelength), we asked the algorithm to predict the intensities needed for each LED channel to achieve the measurements. These predicted intensities were then compared to the original (true) intensities that were used (i.e. predicted - true intensity) as a measure of the accuracy of the prediction for each LED channel. Mean squared error (MSE) was used to summarise the accuracy of the predictions across all LED channels for a given event. To benchmark our algorithm, we compared our multidimensional approach (either with NNLS or SLE) with three other approaches, which treat each LED channel individually: (i) taking the closest intensity from the calibration data [32]; (ii) carrying out a separate linear regression across each LED channel [31]; and (iii) computing a non-negative least squares regression across each channel individually.

We observe that our multidimensional approaches consistently show lower MSEs compared to the individual approaches (Fig. 4A) (p<0.01, Table S1), with a particular advantage as the complexity (i.e. number of LED channels active) increases. We note that multidimensional SLE and multidimensional NNLS produce very similar MSEs for this test, with some differentiation with outliers.

**Figure 4.**
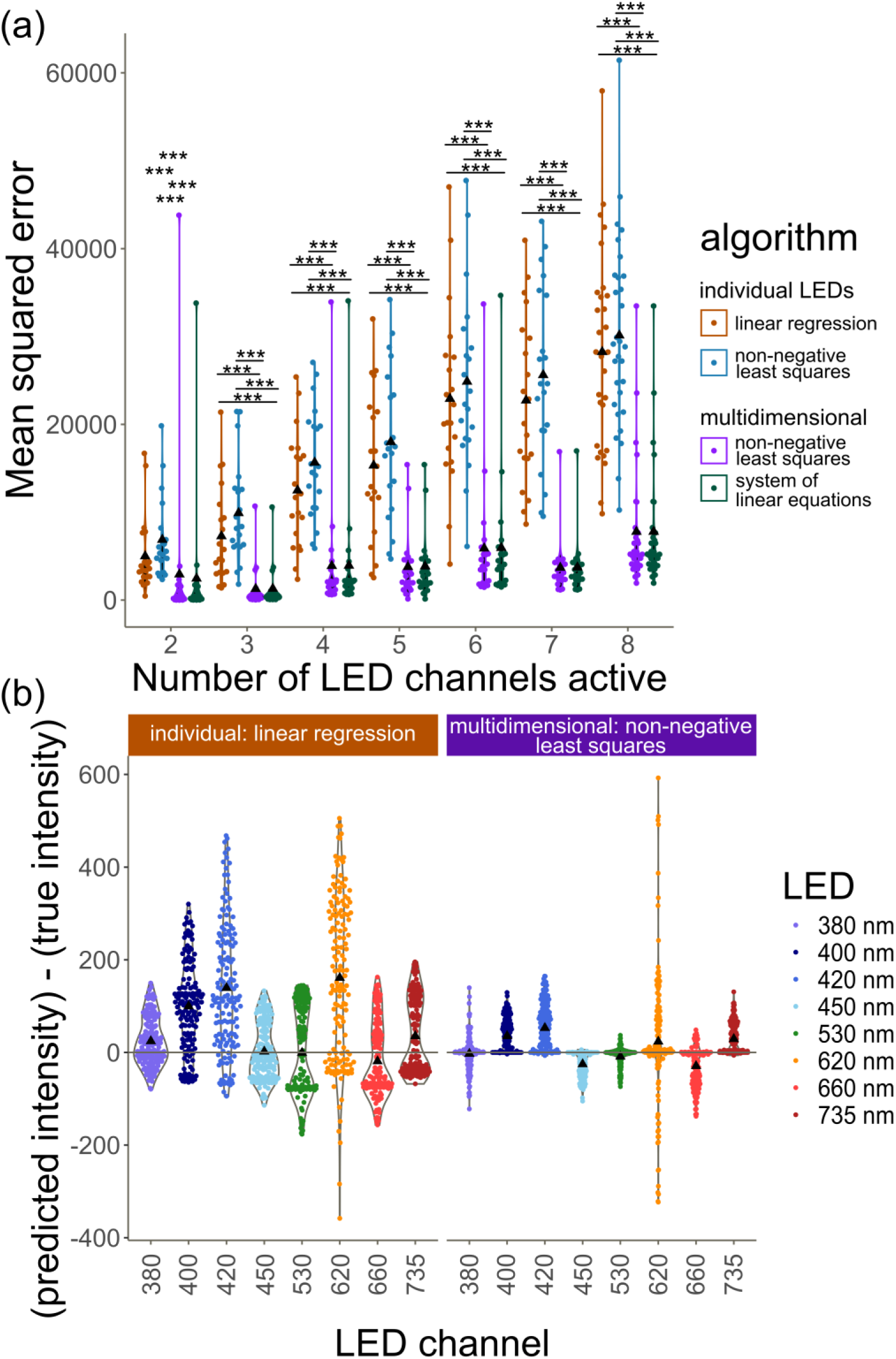
Testing our algorithm against previous methods. The algorithms were tested on their ability to predict the correct intensities which were originally used in a light regime with random intensities in random combinations of LEDs at increasing complexity. Individual LED approaches (closest, linear regression, non-negative least squares) were compared with multidimensional methods (system of linear equations, non-negative least squares). ▲ denotes the mean. **(a)** Mean squared error of each event at increasing complexities. For all algorithms, see figure S2. (***) indicates p<0.001 in pairwise comparisons within the complexity level. **(b)** Residuals ([predicted intensity] - [true intensity] i.e. how wrong the prediction is) per LED across all complexity levels. The best individual approach is compared with a multidimensional approach to illustrate the differences. For the LED breakdowns across all algorithms, see **figure S3**.

We investigated whether the final tidying step of our algorithm had an effect on the accuracy and found that the pre-tidied (i.e. raw predicted values) predictions did not have a substantial difference in MSE (Fig. S2) compared with tidied predictions.

When comparing linear regression (the best of the individual approaches) and multidimensional NNLS (the best multidimensional approach) (Fig. 4B), we see that both consistently predict higher intensities than the true intensity in the blue (380, 400, 420, 450 nm) channels but this effect is markedly less pronounced with the multidimensional approach. We hypothesise that this is due to our approach being able to account for the interaction between LED channels, particularly in the blue range (B), where there is significant bleedthrough from overlapping spectra.

We note that the near-red (620 nm) channel performs consistently poorly across all algorithms. We hypothesise that this is due to the non-monotonic relation between intensity and irradiance (Fig. 3A).

### Refinement using random search

To assess whether our algorithm’s predictions could be improved upon, we carried out an additional refinement process using a random search. Here, intensities (within a defined range) are randomly selected in an attempt to generate an irradiance closer to the target (Fig. S4A). We selected target irradiances where our algorithm predicted intensities generating low, medium and high overall MSEs in order to assess the marginal benefit of this additional refinement.

Regardless of the MSE of the algorithm predicted intensities (i.e. the starting MSE), the random search was able to improve the accuracy of the lights (Fig. 5). The high starting MSE seems to have benefited most from this refinement process, whereas users may not need to invest extra time in refinement if the intensities predicted by the algorithm are already very close to the target.

**Figure 5.**
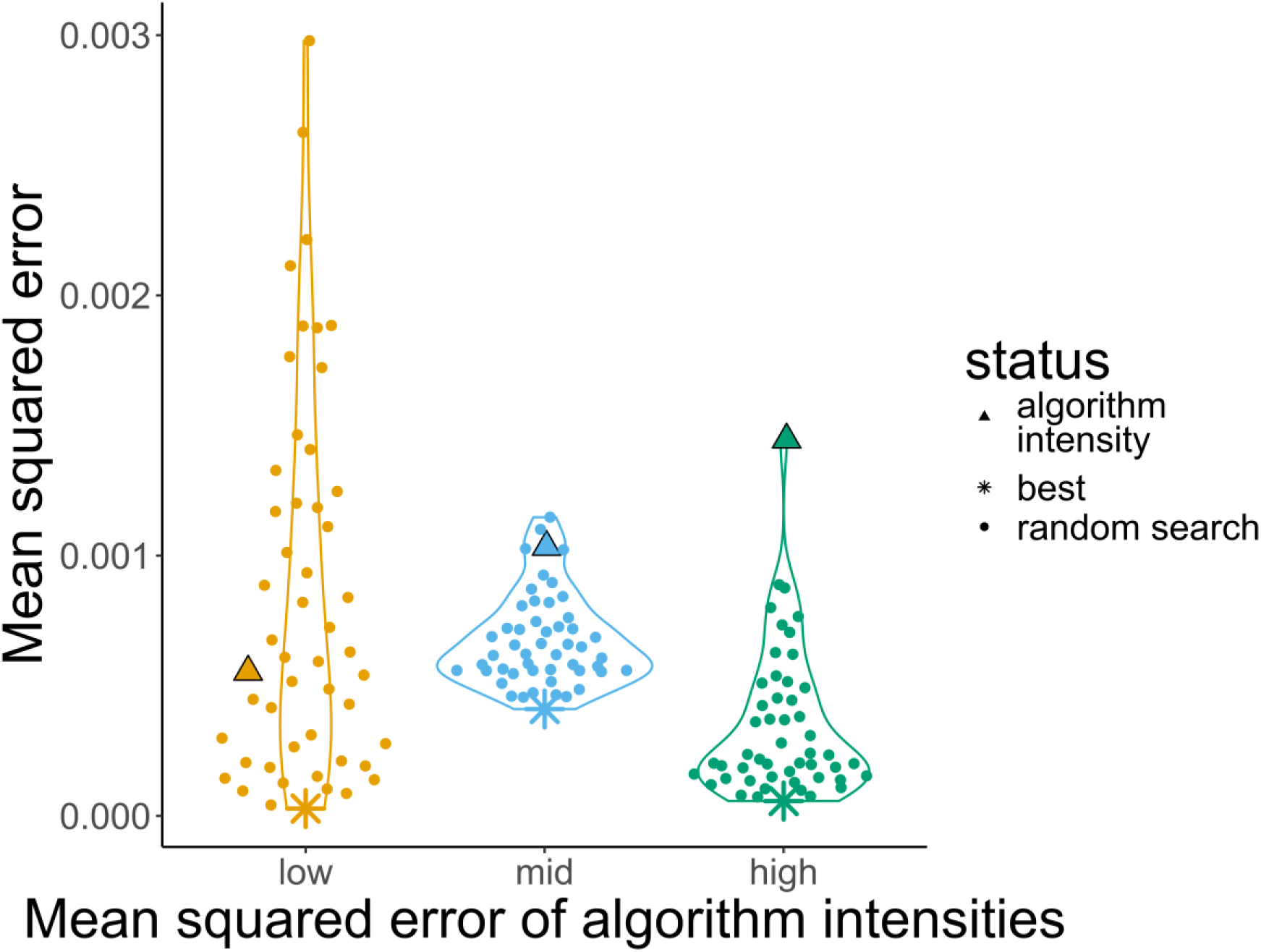
Random search. Target irradiances (complexity: 4 LEDs) were chosen based on the accuracy of the intensities predicted by the multidimensional NNLS algorithm, with a low, medium and high mean squared error (of the algorithm prediction) selected (▲). These algorithm-predicted intensities were used as the basis of a random search in an attempt to find the light intensities with the lowest mean squared error (*). Each point represents an event where random intensities (within a range) were selected for each of the 4 LED channels.

We note that the random search for the mid starting MSE treatment was not able to reduce the MSE as much as the low and high treatments. We hypothesise that the challenge in identifying intensities in this treatment comes from the inclusion of the 620 nm channel (Fig. S4A), which as previously noted had a non-monotonic relation between intensity and irradiance (Fig. 3A).

When taking measurements using any instrument, there is an amount of random technical noise which affects the accuracy of the measurement. Since the MSEs we observe are quite small, we wished to determine whether the improved accuracy from the random search arose because we were truly approaching the local optimum intensity rather than capturing random noise in spectrometer readings. To assess this, we determined whether the LED intensity combinations that were more similar (i.e. small Euclidean distance) to our final selected intensities produced more accurate irradiances (i.e. lower MSE) than intensities that were further away. If our results are due to random technical noise, we would expect a random distribution of points in Fig.S4b. However, in the mid and high algorithm MSE treatments, Fig. S4B shows a smooth gradient in MSE with a strong positive correlation (Table S2), therefore confirming that we are approaching the local optimum for those treatments and that random search is an effective strategy for refining selected intensities.

## Discussion

Most research on plant light response has been done under discrete conditions, which are dissimilar to the dynamic natural light conditions that they have evolved under [22,35]. Advances in LED technology allow us to program more realistic light regimes which mimic the spectrum and fluctuations of natural daylight at different latitudes and times of year. Indeed, more realistic conditions are beginning to be explored, with implications for fluctuating light affecting leaf morphology and photosynthetic acclimation [24]; natural light resulting in differences in metabolite use [11]; and twilight length affecting growth and flowering [23]. Taken together, these studies show that conclusions from discrete lighting experiments cannot be directly translated to natural conditions. Therefore, dynamic lighting conditions are potentially a fruitful area for further research.

However, the complexity of the light regimes that are possible to investigate is limited by the difficulties of accurately programming LEDs across multiple channels. The algorithm we present here allows us to more accurately programme LED fixtures, thus unlocking the potential of these LEDs for research. By using a multi-step approach, LightFitR (https://github.com/ginavong/LightFitR) can account for a non-linear relationship between intensity and irradiance output, as well as channels with overlapping wavelength ranges. Our approach is light-agnostic, meaning that it can work for any programmable LED-based lights which have multiple channels. This package therefore allows the community to more accurately program complex light regimes involving multiple LED waveband channels, fine-tuning wavelength ratios as well as gradual changes to irradiance through the day.

We have shown the value of calibrating the lights when installing them in a cabinet, and [36] suggests calibrating them regularly thereafter. Although some LEDs in some scenarios show more linearity between intensity and irradiance [31], this is not the case universally (Fig 3A). Further, we obtained the full spectrum for each channel, providing insight into bleedthrough between the waveband channels. Thus, in addition to calibration, we encourage plant scientists to more comprehensively report their lighting conditions in controlled environment experiments, in line with [30].

We have demonstrated that our multidimensional algorithm outperforms other approaches, especially at high complexities. Other approaches assume linearity between intensity and irradiance [31], and treat each LED channel as an independent variable. Our approach accounts for potential non-linearity by first subsetting to the calibration point which is closest to the target. Then by using a multidimensional system of linear equations, we can account for the bleedthrough between different LED channels. This may not be required if the number of active LED channels (i.e. complexity) is low. However, as the research is moving towards more complex regimes which emulate natural / field conditions, there will be a need for methods of reliably programming them.

Additionally, we present an extra refinement step which uses a local random search. This represents a way to provide additional accuracy for light regimes that require it, and is more efficient than previous measure and adjust approaches [32]. However, it is a rather time consuming step and so we recommend it only if the extra accuracy is needed and existing methods are not getting users close enough to the target irradiances. Though not demonstrated here, a more thorough grid search, where every possible combination of intensities is tested within a narrow range, can be used to further refine the random search. However, the time required to do grid search increases exponentially with the number of LED channels involved. Therefore we would suggest doing grid search only if absolutely necessary.

The methods presented here provide a way for the plant sciences community to accurately program complex light regimes on LED-based lights. This will allow researchers to investigate more natural regimes, where irradiance and wavelengths change gradually through the day. This could lead to deeper insights into, for example, how variation in light quality and quantity entrains the circadian clock; how plants respond to temporary shade (e.g. from clouds or upper canopy); and how seasonal changes in light quality affect phenological responses. Our approach may also be beneficial for applied research, for example optimising light regimes for indoor controlled environment agriculture and vertical farms. We look forward to seeing how the community uses our package.

## Conclusions

Here, we present a method for accurately programming complex light regimes using multiple LED channels. By using a multi-step, multidimensional algorithm, we can account for non-linearity between intensity settings and light irradiance output as well as bleedthrough between different LED channels, outperforming other approaches. The algorithm-predicted intensities can be further refined using a localised random search. This approach will enable the plant sciences community to explore plants’ responses to complex light regimes, such as the conditions experienced in the field.

## Declarations

## Acknowledgements

We thank the Horticulture Team at the University of York Biology Department for their maintenance of the growth facilities, cabinets and helpful insights. We are also grateful to David Nelmes for IT support with the lights, and Ethan Redmond for stimulating discussion about the mathematics used in this paper.

## Author contributions

GV and DE designed the algorithm and wrote the R package. PS and JD installed Heliospectra lights into the growth cabinets, and made additional modifications to the cabinet. GV, WC, PS and JD measured and tested the lights. GV carried out the experiment design, analysis and figure creation. KD and DE supervised the work. All authors contributed to the writing of this manuscript.

## Data & software availability

The R package (LightFitR) containing the algorithm is available under the GNU General Public License and can be installed from GitHub: https://github.com/ginavong/LightFitR

The processed data and analysis code (in R) for this paper is available under the GNU General Public License in the 2024_LightFitR_MethodsPaper GitHub repository: https://github.com/ginavong/2024_LightFitR_MethodsPaper

The raw datasets supporting the conclusions of this article are available on Zenodo: https://doi.org/10.5281/zenodo.15584172

## Ethics approval and consent to participate

Not applicable

## Consent for publication

Not applicable

