## Supplementary Figures for "Accurately Programming Complex Light Regimes with Multi-channel LEDs"

### Supplemental Figures

## S1

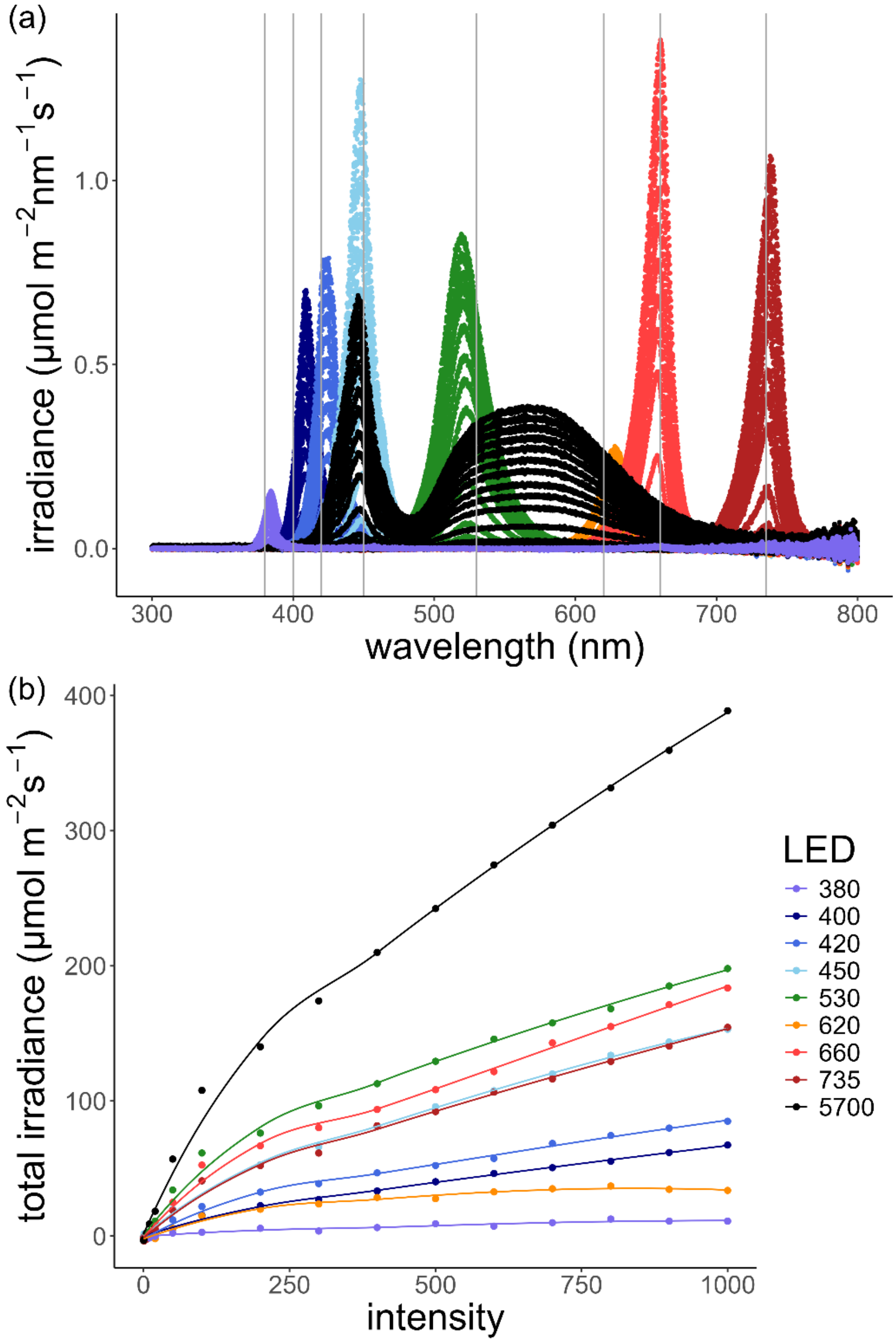


**Figure S1.** Calibrating the lights. **(a)** Spectrum produced by LEDs as each channel was calibrated at increasing intensities. The vertical lines represent the purported wavelengths. **(b)** Total irradiance (across the spectrum) at each intensity.

# S2

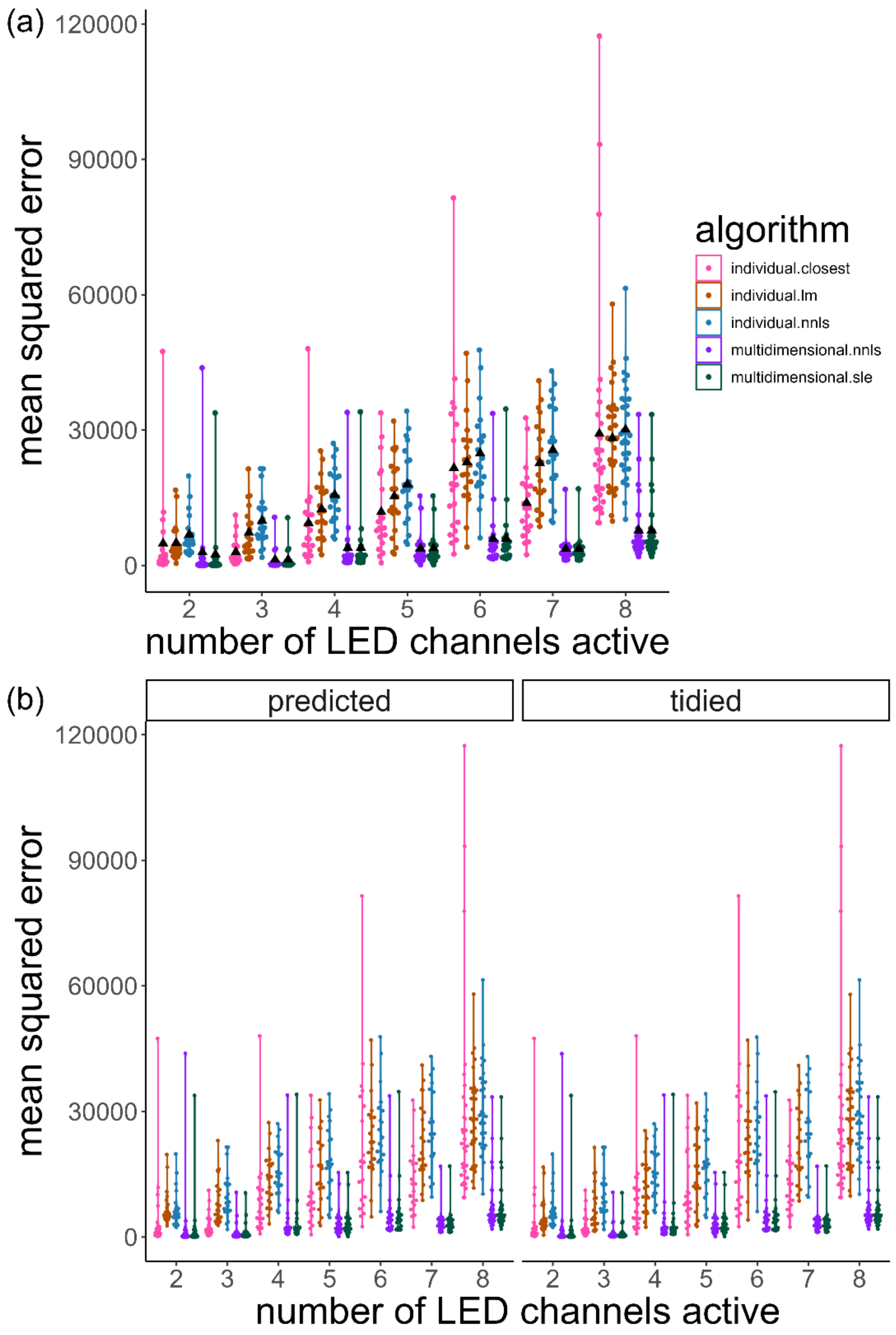


**Figure S2.** Algorithm comparisons. ▲ denotes the mean. **(a)** Mean squared error of indivIdual and multidimensional algorithms at different complexity levels. **(b)** Mean squared error of the raw predictions of the various algorithms vs after the final tidying step.

# S3

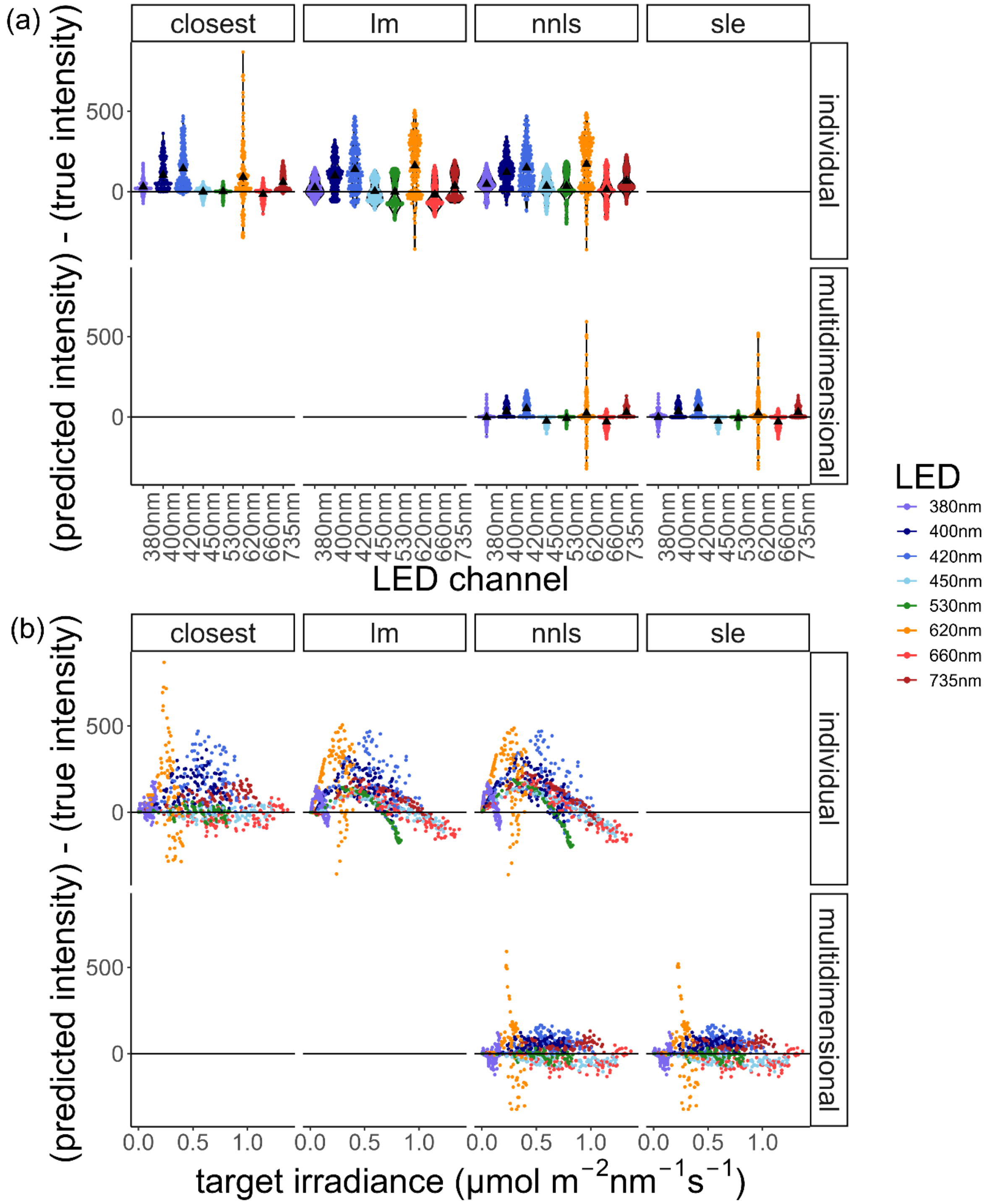


**Figure S3.** Comparing the performance of the algorithms at the LED level. **(a)** The distribution of residuals for each of the algorithms. ▲denotes mean. **(b)** The relationship between residuals and target irradiance.

# S4

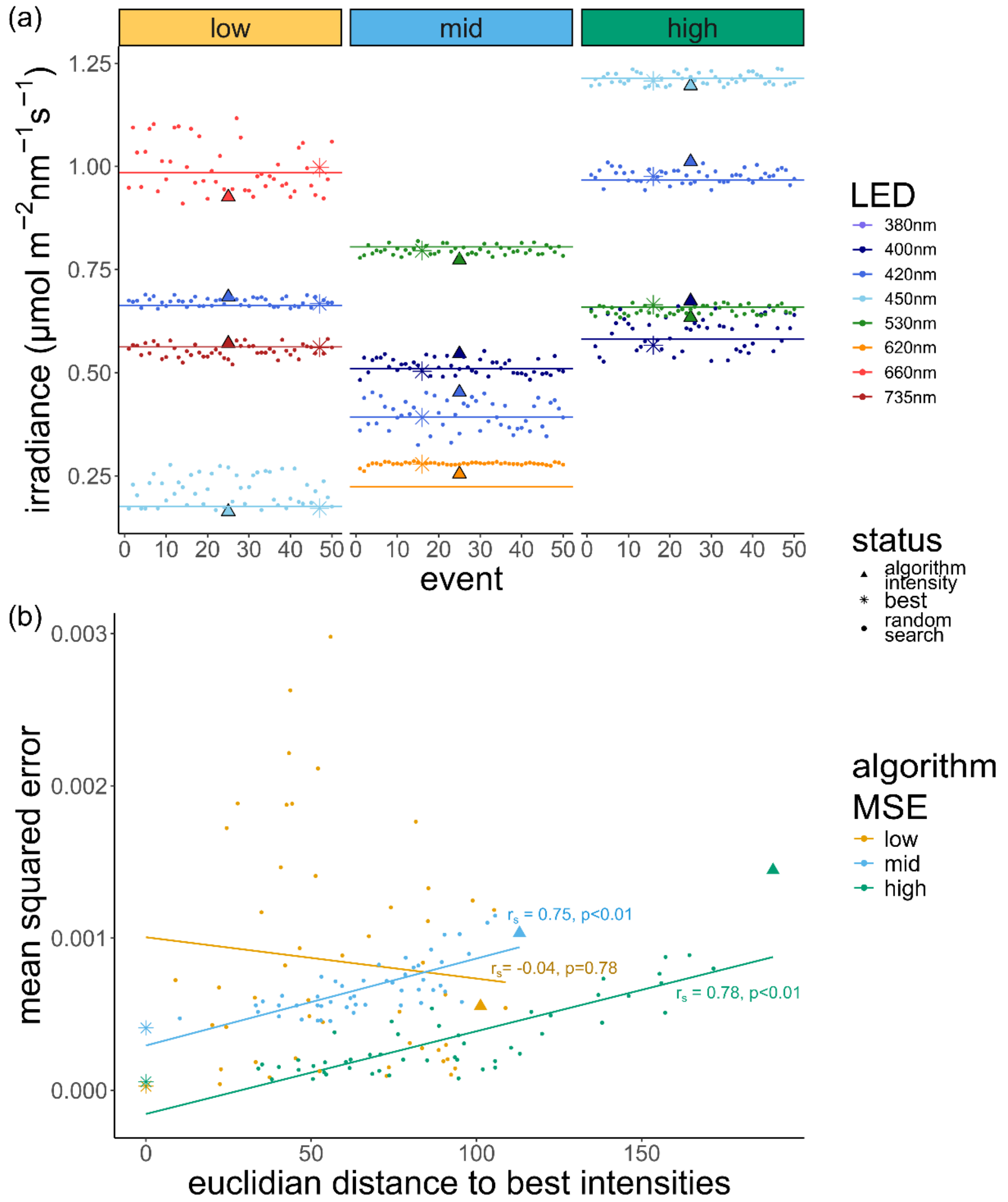


Figure S4. Random search. ▲ is the starting point (i.e. intensities predicted by the multidimensional NNLS), * is the best intensity (i.e. lowest mean squared error) found through random search. **(a)** The progression of the random search for each of the starting MSEs (low, medium high). The horizontal line (-) marks the target irradiance for each of the LEDs. **(b)** The relationship between the straight line distance (i.e. euclidian distance) to best intensities (across all LEDs involved), and the mean squared error. Each point represents an event, with a linear regression (solid line, -) and Spearman rank correlation coefficient (rs) for each starting MSE.
